# Novel engineered chimeric engulfment receptors trigger T-cell effector functions against SIV infected CD4+ T cells

**DOI:** 10.1101/2022.06.21.495546

**Authors:** Daniel Corey, Francoise Haeseleer, Lawrence Corey

**Affiliations:** CERo Therapeutics, San Francisco, CA, U.S.A.; Department of Laboratory Medicine, University of Washington, Seattle, WA, U.S.A.; Vaccine and Infectious Disease Division, Fred Hutchinson Cancer Research Center, Seattle, WA, U.S.A.

**Author notes:** Correspondence should be addressed to D.C. Daniel Corey, 201 Haskins Way, Suite 230, South San Francisco, CA, 94080. Outpace Bio, Seattle, WA, 98109 USA. Authors made equal contribution (Daniel Corey, Francoise Haeseleer).

**Keywords:** Chimeric engulfment receptor, SIV, HIV, CER, TIM-4, phosphatidylserine

## Abstract

Adoptive therapy with genetically engineered T cells offers potential for infectious disease treatment in immunocompromised persons. HIV/simian immunodeficiency virus (SIV) infected cells express phosphatidylserine (PS) early post-infection. We tested whether chimeric engulfment receptor (CER) T cells designed to recognize PS-expressing cells could eliminate SIV infected cells. Lentiviral CER constructs comprised of the extracellular domain of T-cell immunoglobulin and mucin domain containing 4 (TIM-4), the PS receptor, and engulfment signaling domains were transduced into primary rhesus macaque (RM) T cells. We measured PS binding and T-cell engulfment of RM CD4+ T cells infected with SIV expressing GFP. As chimeric antigen receptor (CAR) T cells induce PS and subsequent TIM-4 binding, we evaluated in vitro killing of CAR and CER T-cell combinations. We found that recombinant TIM-4 bound to SIV infected cells. In vitro, CER CD4+ T cells effectively killed SIV infected cells, which was dependent on TIM-4 binding to PS. Enhanced killing of SIV infected CD4+ T cells by CER and CAR T-cell combinations was observed. This installation of innate immune functions into T cells presents an opportunity to enhance elimination of SIV infected cells and offers potential to augment functional cure of SIV/HIV infection.

## INTRODUCTION

Cell therapies are being investigated to treat a large variety of diseases. Chimeric antigen receptor (CAR) T cells are cells engineered to recognize and kill clinically relevant targets. CAR T-cell therapies are an example of successful cell therapy, especially to treat different types of hematologic malignancies^1, 2^. New cells therapies are being investigated to broaden the number of diseases where they might be applicable and explore the potential of other effector cell types such as macrophages and natural killer cells and other CAR^3-6^. The application of the knowledge resulting from in vivo studies and clinical trials with CAR T cells could lead to more successful cell therapies targeting HIV reservoirs that remain in HIV-infected patients and in controlling viral rebound^7-9^.

A common feature displayed by cells that have become apoptotic because of viral or parasitic infections, aging, or altered metabolism is the redistribution of phosphatidylserine (PS) to the outer leaflet of their plasma membrane^10^. For example, HIV infection was shown to trigger the exposure of PS by activating scramblases^11^. The exposed PS was also shown to facilitate fusion between the viral and cell membrane^11, 12^. In addition, because HIV viruses are produced by apoptotic CD4 T cells, HIV virions contain PS in their envelope that stimulates clearance mechanisms and facilitate entry into other cell types such as macrophages^12, 13^.

The exposure of PS on the surface of apoptotic cells is a key “eat me” signal triggering engulfment by phagocytes^10^. Several PS receptors have been identified, including the T-cell immunoglobulin and mucin domain containing 4 (TIM-4) receptor, and anti-TIM-4 antibodies can block this engulfment process by macrophages^14, 15^.

In this study, we developed novel chimeric engulfment receptors (CER) that take advantage of TIM-4 binding PS ^16^. These CERs are composed of the extracellular TIM-4 domain and one or more intracellular signaling domains from receptors involved in innate immune responses to pathogens; e.g., Toll-like receptors (TLR) ^17^. Addition of innate immune function such as phagocytosis, antigen presentation, and greater lytic and non-cytolytic killing offers the potential of enhancing immune responses to chronic HIV infection. These CERs expressed in T cells provided the capability of eliminating simian immunodeficiency virus (SIV) infected cells in vitro. This investigation provides the initial rationale for use of CER T cells in in vivo models of nonhuman primate lentiviral infection to determine if the addition of functional killing and other innate functions such as enhanced antigen presentation and reversal of endogenous T helper responses can be improved through adoptive transfer experiments of T cells with enhanced engulfment function.

## RESULTS

### TIM-4 binds to SIV infected CD4+ T cells

Exposure of PS occurs when HIV infects CD4+ T cells ^11^. To visualize TIM-4 binding to PS exposed on the surface of SIV infected cells, we created a fluorescent SIVmac239 virus ^18^. The coding sequence for the enhanced green fluorescent protein (EGFP) was introduced between the matrix and capsid domains of the SIVmac239 Gag protein and flanked by SIV protease cleavage sites (SIVGAGGFP) (**Figure S1**).

To test if TIM-4 bound to SIV infected cells, we used a TIM-4 Fc chimera composed of the TIM-4 extracellular domain fused to the N-terminus of the human IgG Fc region. When CD4+ T cells were infected with SIVGAGGFP, strong binding of TIM-4 Fc to SIVGAGGFP+ cells was detected 1 hour post infection while no binding was observed on SIVGAGGFP-gated cells (**Figure 1A**). Binding of the labeled anti-IgG antibodies to SIVGAGGFP+ cells was not observed in the absence of TIM-4 incubation, thus confirming interaction of TIM-4 to SIV infected cells (**Figure 1B**). As a positive control, TIM-4 bound to cells undergoing apoptosis triggered by treatment with camptothecin (**Figure 1C**).

**Figure 1.**
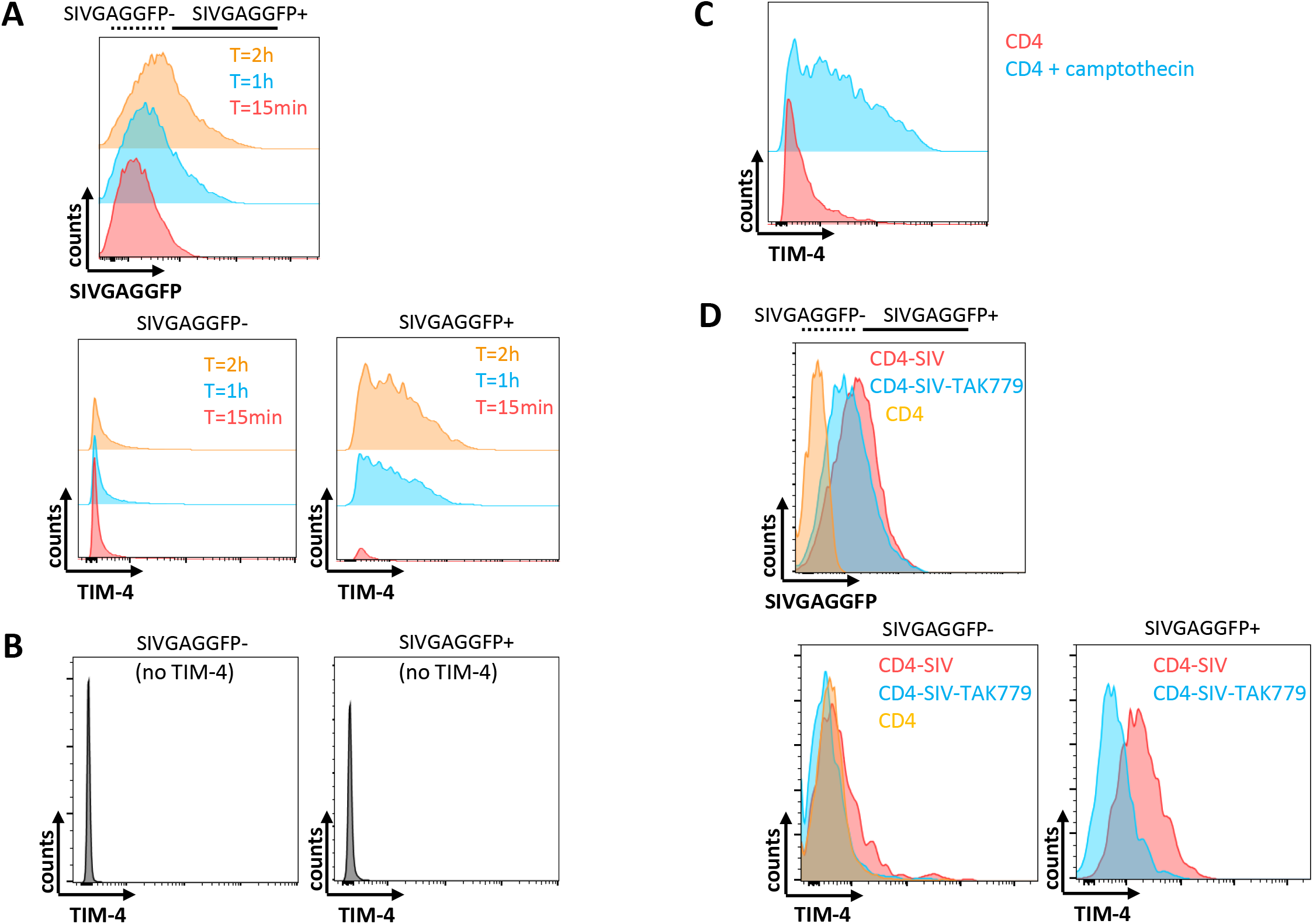
TIM-4 binds to SIVGAGGFP infected CD4+ T cells. A) Flow analysis of recombinant human TIM-4 binding to cells infected with SIVGAGAGFP for 15 minutes (red), 1 hour (blue), or 2 hours (orange). Lower plots show binding when gated on uninfected (SIVGAGGFP-) cells (left) or infected (SIVGAGGFP+) cells (right). B) Same experiment as in A without addition of TIM-4. Uninfected (SIVGAGGFP-) cells (left) or infected (SIVGAGGFP+) cells (right). C) TIM-4 binding to CD4+ T cells incubated in the presence (blue) or absence (red) of 2 μM camptothecin for 24 hours. D) Overlaid flow cytometry histograms of SIVGAGGFP infected CD4+ T cells in the presence (blue) or absence (red) of TAK779; uninfected CD4+ T cells are shown in orange (upper panel). Lower plots show binding when gated on uninfected (SIVGAGGFP-) cells (left) or infected (SIVGAGGFP+) cells (right).

The CCR5 coreceptor is necessary for SIV infection of CD4+ T cells. CCR5 antagonist TAK-779 blocks the interaction between CCR5 and SIV and inhibits PS exposure induced by R5-tropic virions ^11, 19, 20^. Incubation of SIVGAGGFP with CD4+ T cells in the presence of TAK-779 decreased virion binding to target cells and also decreased TIM-4 binding to infected T cells (**Figure 1D**). These results indicate that TIM-4 detects PS exposed on the surface of SIV infected cells.

### CER T cells can kill SIV infected cells upon infection

We initially constructed a series of prototype CER in order to evaluate if differences in signaling domains altered functional activity in vitro. All DNA constructs used the TIM-4 extracellular domain. As TLRs are known to enhance endosomal transfer and trafficking, some of the constructs tested contained a TLR signaling domain (**Figure 2A**). The DNA constructs included a truncated version of the epidermal growth factor receptor (EGFR), which can be detected on the cell surface using an anti-EGFR monoclonal antibody, to assess the efficiency of lentiviral transduction into T cells. We also introduced a membrane anchored fusion inhibitory peptide derived from gp41 (C46) to protect the CD4^+^ CER T cells against SIV infection ^21, 22^. We first investigated if CD4+ or CD8+ CER T cells would be efficient in eliminating SIV infected T cells. Transduction of RM CD4+ and CD8+ T cells with CER21 or EGFR lentivirus led to high levels of EGFR expression (**Figure 2B**). We developed a real-time fluorescence assay to evaluate CER T-cell potency against freshly SIVGFP infected target cells expressing surface-exposed PS ^23^. A significant decrease in the number of GFP+ infected T cells were detected over time in the presence of CD4+ CER T cells but not CD8+ CER T cells or EGFR T cells, indicating the potency of CD4 CER T cells in killing SIV-infected cells (**Figure 2C**). To direct CER T cells to major sites of SIV/HIV persistence, the cDNA encoding RM CXCR5, a homing receptor shown to promote cell trafficking to B-cell follicles in lymph nodes, was added to the lentiviral vector ^24-28^. About 16% of the EGFR+ transduced T cells expressed both EGFR and CXCR5 (**Figure 2D**). CD4+ CER T cells transduced with the CER21-CXCR5 lentivirus were efficient in killing SIV infected targets (**Figure 2E**).

**Figure 2.**
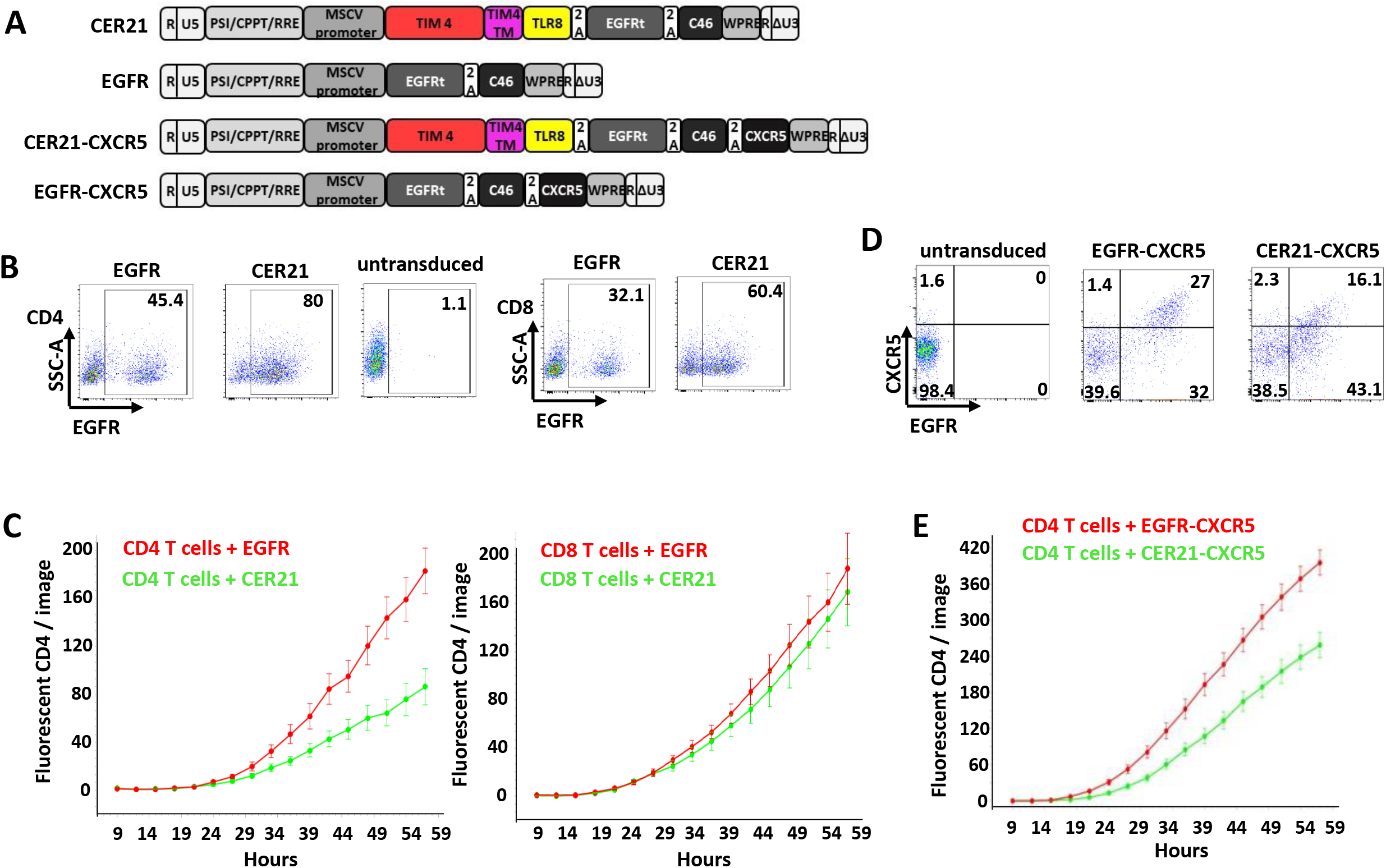
Assessment of CER T-cell potency in killing SIV infected cells. A) Schematic diagram of the CER construct with the extracellular domain of TIM-4 linked to the TLR8 signaling domain (CER21), CER21 with CXCR5 (CER21-CXCR5), the EGFR control vector (EGFR), and EGFR with CXCR5 (EGFR-CXCR5). B) Flow cytometry analysis of CD4+ and CD8+ T cells transduced with either CER21 or EGFR gated on expression of EGFR. Numbers in plots indicate the percentage of EGFR+ cells after gating on lymphocytes and singlets. Control untransduced CD4+ T cells are also shown. C) SIVGFP infected CD4+ T cells were mixed with CER T cells (green) or control EGFR T cells (green) at t=0. The E:T ratio was 5:1 for both CD4 (left) and CD8 (right) assays. Five images were taken in triplicate wells every 3 hours. The error bars indicate the standard error to the mean. D) Flow cytometry analysis of CD4+ T cells transduced with CER21-CXCR5 or EGFR-CXCR5 for expression of EGFR and CXCR5. E) Assessment of CER21-CXCR5 T-cell potency in killing SIV infected cells as described in C.

### CER composed of a TLR8 signaling domain and CD3ζ activation domain is the most potent in killing SIV infected cells

We designed 8 additional CERs by linking the extracellular domain of TIM-4 to various intracellular signaling domains with either the TIM-4 or CD28 transmembrane domain. The intracellular domain was composed of one or multiple engulfment signaling domains of TLR2, TLR8, tumor necrosis factor receptor associated factor 6 (TRAF6), DAP10, DAP12, and/or CD28. Some CERs also included TLR signaling domains together with the CD3ζ activation domain (**Figure 3A**). High expression of the EGFR marker was observed for all 9 CER-transduced RM CD4+ T cells (**Figure 3B**). When comparing the effector functions of these CER T cells in the real-time fluorescence assay, we found diverse killing potency, with CER131 (TLR8-CD3ζ) and CER29 (TRAF6) exhibiting the greatest effector functions towards SIV infected cells (**Figure 3C**).

**Figure 3.**
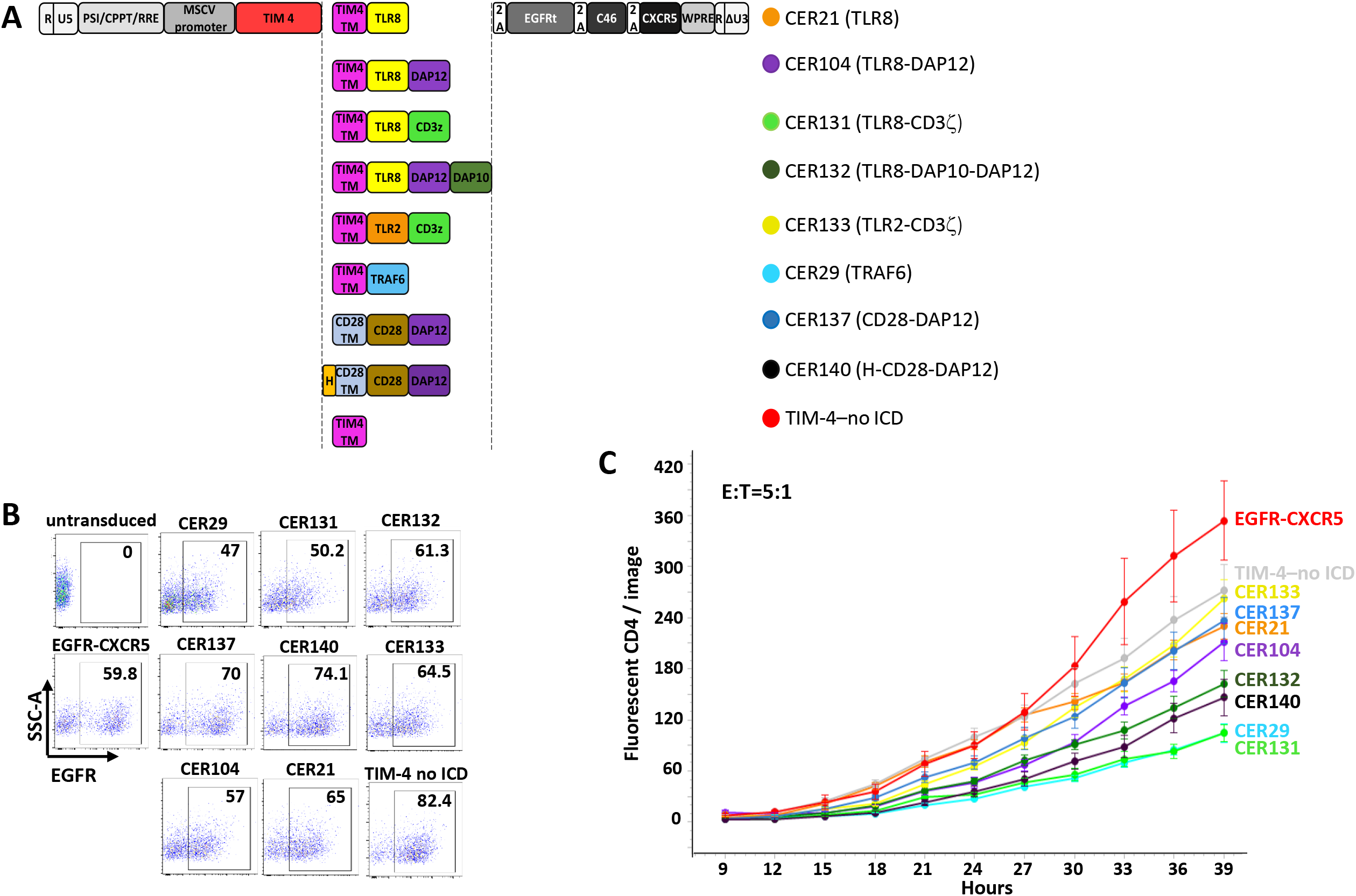
Comparison of effector functions of diverse CERs. A) Schematic diagram of 8 CER constructs with the extracellular domain of TIM-4 linked to one or multiple intracellular signaling domains (ICD) originating from TLR8, TLR2, DAP10, DAP12, TRAF6, CD28 or CD3z; and negative controls without ICD (TIM-4–no ICD) or EGFR-CXCR5. B) CD4+ T cells transduced with CER lentiviruses. C) Assessment of CER T-cell potency in killing SIV infected cells as described. Five images were taken in triplicate wells every 3 hours. The error bars indicate the standard error to the mean. *P* values <0.05 for CER29, CER131, CER140, CER132 and CER21 T cells compared with EGFR-CXCR5 T cells (Student’s t-test). *P* values <0.05 for CER29, CER131, CER140 and CER132 T cells compared with TIM-4-no ICD T cells.

### Mutations in the PS binding site of TIM-4 impair CER T-cell effector function

In the immunoglobulin domain of TIM-4, a hydrophobic phenylalanine-glycine (FG) loop is located in a cavity important for metal ion interaction and PS binding ^29^. Mutations of 4 amino acids, tryptophan, phenylalanine, asparagine and aspartic acid (WFND), present in this cavity result in loss of phagocytosis of apoptotic cells ^14^. In order to determine whether CER effector functions are triggered upon specific recognition of PS by TIM-4, we generated CER constructs with alanine mutations or deletion of all 4 WFND residues (**Figure 4A**). After assessment of the transduction rate of CER T cells (**Figure 4B**), CER mutants were tested in the real-time killing assay. Both mutants were impaired in their ability to eliminate SIV infected CD4+ T cells compared with wild-type CER131 (**Figure 4C**).

**Figure 4.**
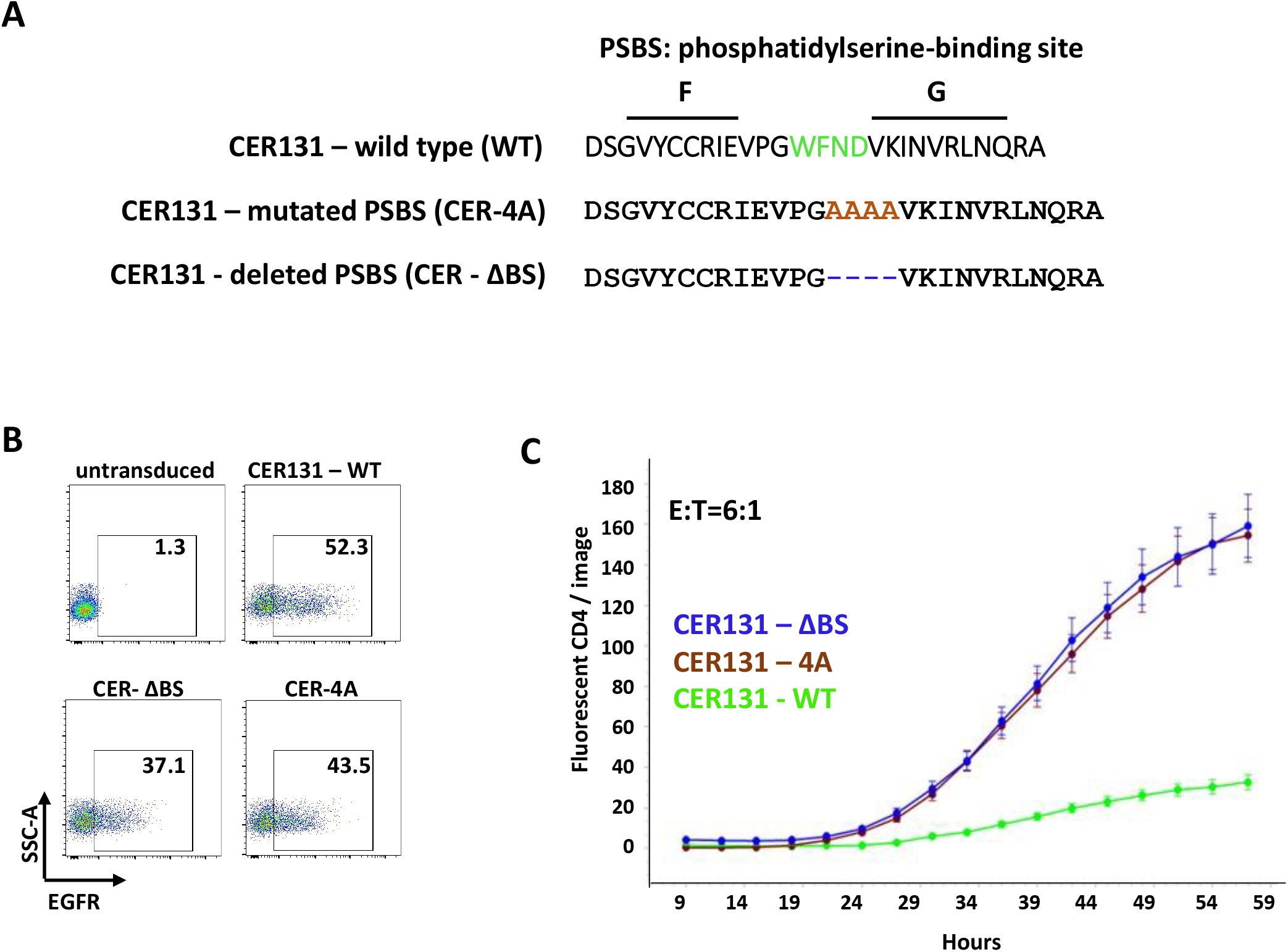
CER131 containing TIM-4 PS binding site (PSBS) mutations lose ability to kill SIV infected CD4+ T cells. A) Amino acid sequence of the wild-type TIM-4 FG loop including four residues important for PS binding (green). Residues replaced by alanine (brown) in CER-4A or deleted (blue) in CER–ΔBS. B) Flow cytometry analysis of CD4+ T cells transduced with CER131 WT and CER131 mutants for EGFR expression. C) Comparison of the potency of CD4+ T cells expressing CER131 WT (green), CER-4A (black) and CER–ΔBS (blue) in killing SIV infected cells as described in Figure 2C. The E:T ratio was 6:1.

### Additive killing activity of CER T cells and anti-SIV CAR T cells against SIV infected cells

As cytotoxic T-cell killing has been shown to elicit surface exposed PS on target cells, we evaluated potential additive effects between CER T cells and CAR T cells directed at SIV infected CD4+ T cells. For these experiments, we utilized two previously constructed lentivirus directed CAR T cells ^30^. The first, ITS06, contains an scFv directed at the V1 region of SIV envelope and the second, VRC26, contains a V2 loop-directed scFv, which cross-reacts with HIV-1 in vitro and is representative of lower avidity but greater breadth. We evaluated combinations of CD4+ CER T cells with CD8+ and/or CD3+ anti-SIV CAR T cells in killing potency against SIV infected CD4+ RM cells. CD4+ CER T cells co-incubated with CD8+ ITS06 CAR T cells induced additive killing of SIV infected target cells (**Figure 5A**). A dose response cytotoxic effect was observed using a E:T ratio of 5:1 for CER T cells and various E:T ratios of ITS06 CAR T cells; most readily seen at a low E:T ratio of 1:20 CAR T cells. A similar concentration dependent effect was seen in experiments using a combination of VRC26 CAR T cells with CER T cells (**Figure 5B**). These latter experiments used a 40-fold higher concentration of the VRC26 CAR T cells (E:T of 2:1) due to its reduced potency compared to ITS06 CAR T cells. These data suggest additive in vitro killing between CER and CAR T cells.

**Figure 5.**
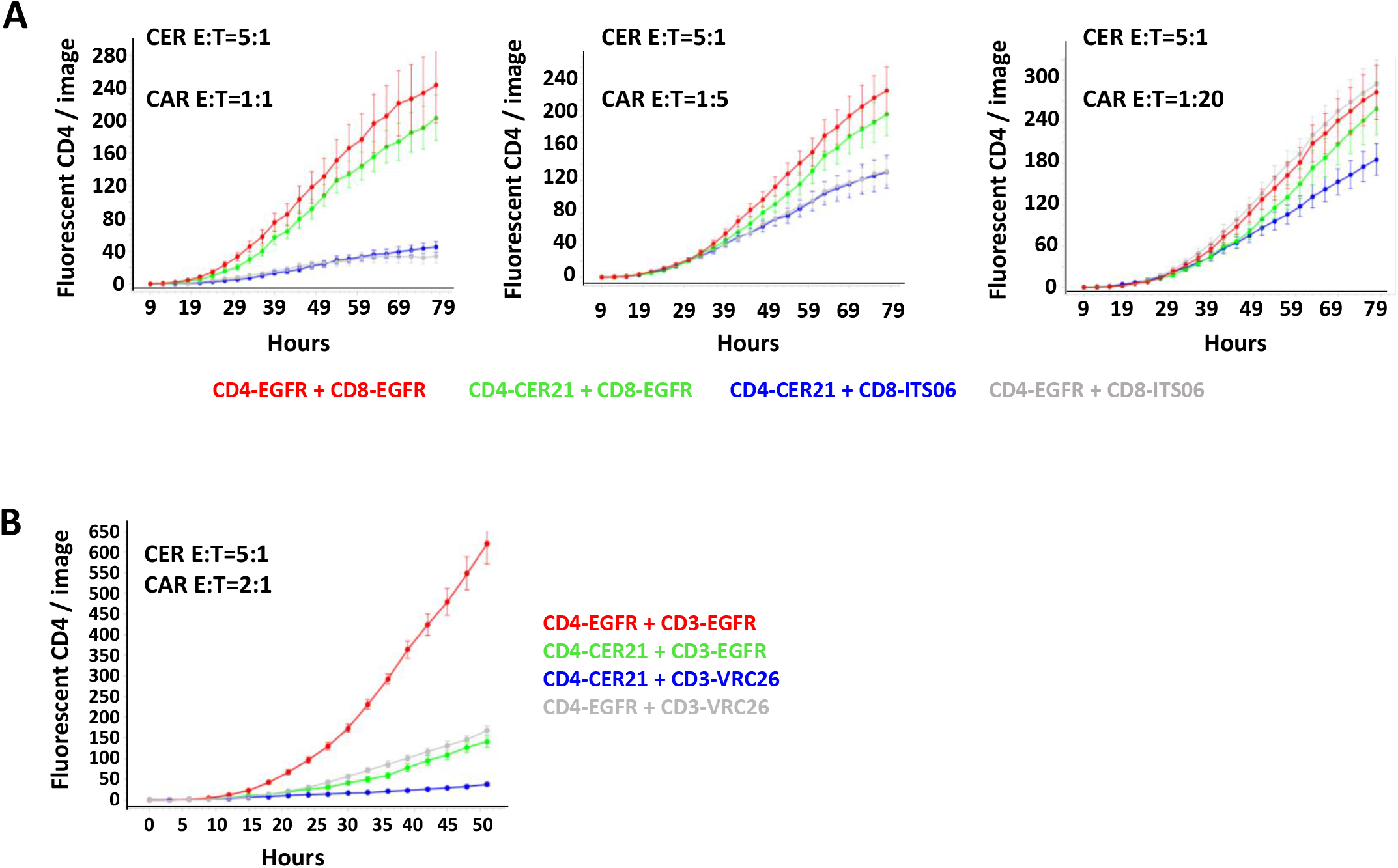
Additive killing activity of CER T cells and SIV directed CAR T cells against SIV infected CD4+ T cells. A) Killing of SIV infected RM T cells by different ratios of CD4 CER and CD8 CAR T cells: EGFR CER with EGFR CAR T cells (red), CER21 with EGFR CAR T cells (green), CER21 with ITS06 CAR T cells (blue), and EGFR CER with ITS06 CAR T cells (gray) at E:T ratios indicated. Five images were taken in triplicate wells every 3 hours for all time points. B) Killing of SIV infected RM CD4+ T cells by the combination of CD4+ CER21 T cells and VRC26 CD3+ CAR T cells.

## DISCUSSION

Although CAR T-cell therapies have shown some efficacy in controlling viral replication, anti-HIV/SIV CAR T cells have not reached the potential shown by CAR T cells targeting cancer cells. Some success in delaying and reducing SHIV or HIV viremia has been primarily achieved with CAR T cells based on the extracellular domain of the human CD4 receptor, while T cells expressing CARs engineered with the scFv of broadly neutralizing antibodies were not successful likely because of anti-SIV antibodies blocking the interaction between the CAR scFv and its epitope ^22, 31-33^. As such, development of novel chimeric receptors targeting different surface proteins and signaling through different immune pathways might lead to improved elimination of HIV infected cells.

Here, we investigated the potential of receptors triggering engulfment signals for the clearance of SIV infected cells. Phagocytes sense PS, the eat me signal, exposed at the surface of apoptotic T cells such as cancer or infected cells, which leads to phagocytic engulfment ^10^. Although multiple PS receptors have been identified, TIM-4 has been shown to be necessary for apoptotic cell engulfment by macrophages ^14, 15^. In this study, we demonstrated that TIM-4 can bind PS exposed on the surface of SIV infected cells and is thus a good candidate for engineering new receptors against SIV.

The TIM-4 PS binding protein in combination with other apoptotic cell clearance pathways led to a range of increased killing by the CER T cells. TLRs, a family of pattern recognition receptor proteins that recognize pathogen-associated molecular patterns and lead to activation of immune signaling pathways, have also been used to boost the immune response against cancer cells and tested as part of CAR T cells ^6, 34, 35^. The TLR8 signaling pathway, including MyD88 and TRAF6, results in NF-κB expression and protection against RNA viruses ^36^. DAP10 and DAP12 are involved in kinase signaling cascades for a variety of immune response receptors ^37, 38^. The CD28 costimulatory domain together with the CD3ζ domain of the T-cell receptor triggers T-cell activation upon antigen recognition, and has been extensively used as signaling component of CAR T cells to promote cytokine production and cytolysis ^39, 40^. We found that out of all the innate signaling constructs we tested, the TLR8-CD3ζ combination and TRAF6 CER T cells killed target cells most efficiently. PS binding by TIM-4 on the surface of CER T cells was essential for killing and mutation in the PS binding pocket abolished activity.

The process triggered by the CER T cells to eliminate SIV infected cells has not been investigated. Prior studies suggest that killing by T cells engineered to express chimeric antigen receptors for phagocytosis (CAR-Ps) appears to be related to the cell’s ability to nibble plasma membrane fragments of other target cells (i.e., trogocytosis) ^3, 41-43^. CERs with diverse signaling domains provide new functionality to CD4+ T cells, and it is possible that different processes are triggered by each receptor ^16^. For example, CER131 that includes a CD3ζ activation domain might also trigger the activation of natural T-cell effector functions. Although no receptors similar to the CERs described in this study have been investigated and tested in T cells, CAR-Ps are another type of engulfment receptor composed of a specific scFv fused to intracellular signaling domains that contain immunoreceptor tyrosine-based activation motifs. These CAR-Ps when expressed in macrophages induce engulfment of specific targets, including cancer cells ^3, 44^. However, for cell therapy, gene transfer into primary macrophages as well as manufacturing primary macrophages for infusion would likely be more complex and expensive than for T cells.

While our data showing such an approach to eliminate SIV infected cells are provocative, several limitations exist. Whether SIV infected T cells are inhibited and put into endosomal pathways and “directed” are not yet known, although evidence for this has been seen with tumor cell lines ^16^. Similarly, whether elimination is achieved by the CD4+, CD8+ or both populations of T cells remains to be determined. In ongoing experiments, most engulfment activity appears restricted to the CD4+ T-cell populations^45^. While a high safety profile has been shown in small animal models, whether off target effects will occur in NHP remains to be seen^46^.

In conclusion, we engineered novel types of chimeric receptors that provide CD4+ T cells the ability to eliminate SIV infected cells in vitro. These genetically engineered CER T cells enhanced in vitro cellular damage caused by high and low affinity CAR T cells, suggesting the approach could be evaluated in vivo. Lastly, in vivo studies will be required to define if the potency and ability of engulfment to enhance antigen presentation and endogenous immune responses will occur in vivo with limited toxicity. To date, extensive animal model experiments in mice in tumor models have shown no evidence of hematologic or systemic toxicity^45, 46^.

## MATERIALS AND METHODS

### Enrichment of CD4+ and CD8+ RM T lymphocytes

Frozen peripheral blood mononuclear cells (PBMCs) from Indian genetic background RM (*Macaca mulatta*) were obtained from the Oregon National Primate Research Center in accordance with standards of the Center’s Institutional Animal Care and Use Committee and the *National Institutes of Health Guide for the Care and Use of Laboratory Animals*. Immunomagnetic negative selection (Easy Sep NHP, STEMCELL Technologies, Cambridge, MA) was used to enrich in CD4+ or CD8+ T cells from PBMCs that were cultured in X-vivo 15 media (Lonza) supplemented with 10% FBS, 100 U/ml Pen/Strep, 1 x glutamax and 50 IU/ml human recombinant IL-2 (Peprotech, Cranbury, NJ). Enriched CD4+ and CD8+ T cells were activated with ImmunoCult NHP CD2/CD3/CD28 T-cell activator (STEMCELL Technologies) and incubated 3 days at 37°C in humidified 5% CO_2_.

### Production of SIVGAGGFP and SIVGFP and CD4 T-cell infection

To generate the fluorescent SIVmac239 virus (SIVGAGGFP), we introduced cDNA encoding enhanced green fluorescent protein (EGFP) together with protease cleavage sites and restriction sites between the matrix and capsid domains of Gag of the full-length SIV_MAC239_ proviral DNA using PCR and the NEBuilder HiFi DNA Assembly (New England Biolabs, Ipswich, MA) and used the same strategy as Hubner et al to generate the HIV Gag-iGFP ^18^. The junction between MA and EGFP is as follows: 5’-<u>CCATCTAGCGGCAGA</u>*GGAGGAAATTACCCAGTACAACAA<u>ACGCGT</u>*ATGGCTAGCAAGGGC -3’ where the MA coding sequence is underlined, the protease cleavage site is in italics, the *MluI* restriction site is in underlined italics and followed by the EGFP sequence. The junction between EGFP and CA is as follows: 5’-GACGAGCTGTACAAG*<u>TCTAGA</u>GGAGGAAATTACCCAGTACAACAA*<u>ATAGGTGGTAACTAT</u>-3’ where the EGFP coding sequence is followed by the *XbaI* restriction site in underlined italics, the protease cleavage site in italics and the underlined CA coding sequence. Virus production for SIVGAGGFP and SIVGFP, a recombinant SIVmac239 virus with an IRES-EGFP cassette downstream of the *nef* gene illustrated in our previous work, was performed as previously described ^23^. Briefly, HEK293T cells were transfected with 20 μg of the SIVGAGGFP or SIVGFP plasmid using the calcium phosphate method. The fluorescent viral supernatant was collected 48 hours later, cleared by centrifugation, filtered, and concentrated. Stocks of SIVGAGGFP or SIVGFP viruses were titrated using TZM-bl cells as described by Wei et al ^47^. Infection of CD4 targets was performed by adding ∼ 20μl of concentrated fluorescent SIV viruses to 10^5^ CD4 cells (MOI:0.5) or control Jurkat 76 cells ^48^ plated on retronectin-coated 96-well plates followed by spinoculation for 2 hours at 1,200 x g. The SIV-infected cells were assessed for infection using flow cytometry after gating on lymphocytes and single cells.

### TIM-4 binding assays

Binding of TIM-4 to infected cells was tested using a TIM-4 Fc chimera composed of the extracellular domain of TIM-4 fused to the N-terminus of the Fc region of human IgG (Abcam, Waltham, MA). One μg of TIM-4 in PBS containing 1% BSA was incubated on ice for 30 minutes with 10^5^ washed CD4 T cells infected as indicated above with SIVGAGGFP. After washing, the cells were incubated for 15 minutes on ice with an Alexa Fluor 647-anti-IgG antibody (clone M1301G05, Biolegend, San Diego, CA). Control experiments were performed by skipping the incubation with TIM-4. Binding of TIM-4 to SIVGAGGFP infected cells was analyzed by flow cytometry after gating on lymphocytes, single cells and GFP+ cells. When indicated, the CD4 T cells were incubated with 10 μm TAK-779 (Medchemexpress, Monmouth Junction, NJ) for 15 minutes before adding the SIVGAGGFP virus and infection of cells. Control apoptotic cells were prepared by incubating CD4 T cells with 2 μM camptothecin (Sigma-Aldrich, St. Louis, MO) overnight at 37°C.

### Generation of lentiviral transfer plasmids and transduction into RM T cells

All CER constructs were generated by standard PCR cloning techniques and the PCR products were assembled using the NEBuilder HiFi DNA Assembly (New England Biolabs). CER21 was generated by linking the extracellular and transmembrane domain of human TIM-4 (GenBank™ accession number AAH08988.1, residues 1 to 335) with the signaling domain of the human toll-like receptor 8 (TLR8) (GenBank™ accession number AAQ88663.1, residues 849 to 1041). The CER are fused to a truncated EGFR as a marker to identify transduced cells via a Thosea asigna virus 2A (T2A) self-cleavage peptide. The C46 inhibitory peptide preceded by a T2A self-cleavage peptide and a signal peptide and linked through an IgG2 hinge to the membrane spanning domain of CD34 ^49^ was also added as a PCR product to the CER construct. When indicated, a cDNA encoding the rhesus CXCR5 (Sino Biological, GenBank™ accession number XP_001100017.2) preceded by a porcine teschovirus-1 2A (P2A) cleavage site was also added downstream of the C46 to build the CER-CXCR5. Control EGFR and EGFR-CXCR5 were generated by removing the CER from the above construct using the NEBuilder HiFi DNA Assembly (New England Biolabs). CER 104 and CER 131 were generated by adding the human DAP 12 signaling domain (GenBank™ accession number NP_001166985, residues 51 to 102) or the human CD3z activation domain (GenBank™ accession number NP_932170.1, residues 52 to 164), respectively, downstream of the TLR8 domain in CER21. CER132 was constructed by adding the human DAP 10 signaling domain (GenBank™ accession number NP_055081, residues 70-93) downstream of the TLR8 and DAP12 signaling domains in CER104. The CER 133 is composed of the human TLR2 signaling domain (GenBank™ accession number AAY85648, residues 610 to 784) and CD3z activation domain. CER 129 is composed of the signaling domain of the human TNF receptor-associated factor 6 (TRAF6) (GenBank™ accession number AAH31052, residues 1 to 274). CER140 and CER137 were generated by linking the extracellular domain of TIM-4 (GenBank™ accession number AAH08988.1 residues 1 to 314) to the transmembrane domain and signaling domain of human CD28 with or without the CD28 hinge, respectively (GenBank™ accession number NP_006130.1, residues 114 to 220 (with hinge) or residues 153 to 220 (without hinge). These CER or control EGFR constructs were cloned together with a woodchuck hepatitis virus posttranscriptional regulatory element (WPRE) into a SIV-based lentiviral vector (a generous gift from Dr. Nienhuis, St June Children’s Research hospital, Memphis, TN and Dr. Miyazaki, Osaka University, Japan) ^50, 51^. The production of recombinant lentiviruses was performed as previously described ^23^. Briefly, Lenti-XTM 293T cells (Takara Bio) in DMEM media containing 10% FBS and 100 U/ml Pen/Strep were transfected using the standard calcium phosphate method with 15 μg of the CER transfer vector, 6 μg of the pCAG-SIVgprre (*gag/pol* and *rev* responsive element [RRE]), 4 μg of the *rev/tat* expression plasmid pCAG4-RTR-SIV and 3 μg of the pMD2.CocalG (glycoprotein G of the cocal virus) ^52^. After overnight incubation, cells were washed and added fresh media. One and 2 days later, lentivirus-containing media was collected, cleared by centrifugation at 1,000 x g for 5 minutes followed by filtration on a 0.45 μm Millipore filter, and concentrated (50 X) by ultracentrifugation at 74,000xg for 2 hours at 4°C. The lentivirus stocks were titrated by transduction of Jurkat cells cultured in RPMI media supplemented with 10% FBS and 100 U/ml Pen/Strep using spinoculation for 2 hours at 1,200xg. The percentage of transduced Jurkat cells was quantified by flow analysis for EGFR using the anti-EGFR cetuximab mAb (Erbitux, PE-conjugated at Juno Therapeutics, Seattle, WA).

For the transduction of CD4+ or CD8+ T cells, cells were mixed on retronectin (Takara Bio)-coated plates with CER lentivirus at a MOI of ∼ 20 followed by spinoculation for 2 hours at 1,200xg. Cells were washed about 24 hours after transduction and expanded in fresh complete X-vivo 15 media. Four days after transduction, T cells were analyzed for EGFR expression by flow cytometry using PE-anti-EGFR, BV421 anti-CD4 (OKT4, Biolegend) and APC-Cyanine7 anti-CD8 (SK1, Biolegend). EGFR expression was analyzed using FlowJo and sequential gating on lymphocytes, single cells and then CD4+ or CD8+ T cells.

### Real-time monitoring of cell infection to assess CER T-cell potency in eliminating SIV-infected CD4

The targets were prepared at the time of the assay. A master mix was prepared by adding SIVGFP at a MOI of ∼ 0.5 to 4. 10^5^ CD4 T cells/ml. Fifty μl/well of the mix (20,000 CD4 T cells + SIVGFP) were distributed into BioCoat poly-D-lysine coated flat bottom 96-well plate and spinoculated. CER T cells or control cells were then added at the effector:target (E:T ratio) of 5:1 in triplicate wells. Plates were incubated at 37°C in the IncuCyte S3 LiveCell Analysis System (Sartorius) and five images of each well were recorded every 3 hours and analyzed with the IncuCyte image analysis software to determine the number of infected cells becoming fluorescent overtime.

### ITS06 CAR T-cell preparation

The design and assembly of the ITS06 CAR and VRC26 CAR was previously described ^30^. The ITS06 CAR was composed to the ITS06 scFv in a VH-VL orientation and a medium spacer of 119 amino acids linked to a CD28 transmembrane domain, a 4-1BB intracellular costimulatory domain, and a CD3z activation domain and was also fused via a T2A peptide to a truncated EGFR as a marker to identify transduced cells. The VRC26 CAR vector consisted of the scFv of the VRC26.25 bnAb and a short spacer of 12 amino acids linked to a CD28 transmembrane domain, a 4-1BB intracellular costimulatory domain, and a CD3z activation domain and fused to a truncated EGFR via a T2A peptide. Both the ITS06 CAR and the VRC26 CAR were also fused to a T2A-C46-peptide-P2A-CXCR5 cassette as described above for the CER vector. The production of recombinant lentiviruses for the CAR and the transduction of T lymphocytes with the CAR lentiviruses was performed as described above for the preparation of the CER T cells. Transduction efficiency was determined by flow cytometry analysis using PE-anti-EGFR.

### Real-time assay to assess the additive effect of CER T cells and CAR T cells against SIV-infected cells

The real time assay to monitor infection of RM CD4 T cells was performed as described above except that instead of CER T cells, a mix of CER T cells at a fixed ratio (E:T of 5:1) and various amounts of CAR T cells as indicated in the figures were added together to the SIV-infected targets.

### Statistics

Data of the real time assays are presented as the mean ± standard error of the mean. Statistical significance was analyzed by Student’s t test at time indicated in the figure legend and compared to controls.

## ACKNOWLEDGEMENTS

Our studies were funded by the Gilead Foundation and NIH/NIAID (grant UM1AI126623-05). We thank Dr. Arthur Nienhuis (St June Children’s Research hospital, Memphis, TN, US) and Dr. Jun-ichi Miyazaki (Osaka University, Japan) for kindly providing the SIV-based lentiviral vectors ^50, 51^ and Drs. Serge Barcy, Mindy Miner and Karsten Eichholz for suggestions, critiques and editing of the manuscript. DC has filed patents on the development of CER T cells, which are licensed to Cero Therapeutics; he is employed by Cero Therapeutics. LC is on the Scientific Advisory Board of Cero Therapeutics. FH declares no conflicts of interest.

## AUTHOR CONTRIBUTIONS

Co-first authorship is listed alphabetically. DC conceived the idea of using TIM-4 and designed the CER T cells including in vitro screens for engulfment activity for the lead constructs. FH designed and performed all the in vitro experiments and wrote the first draft of the manuscript. LC planned the studies and collaboration. FH and LC had unrestricted access to all data. All authors agreed to submit the manuscript, read and approved the final draft and take full responsibility of its content.

## Supplemental Material

**Supplemental Figure 1. Generation of a fluorescent SIVmac239 virus**.

