## Supplemental Figure 1 for "Novel engineered chimeric engulfment receptors trigger T-cell effector functions against SIV infected CD4+ T cells"

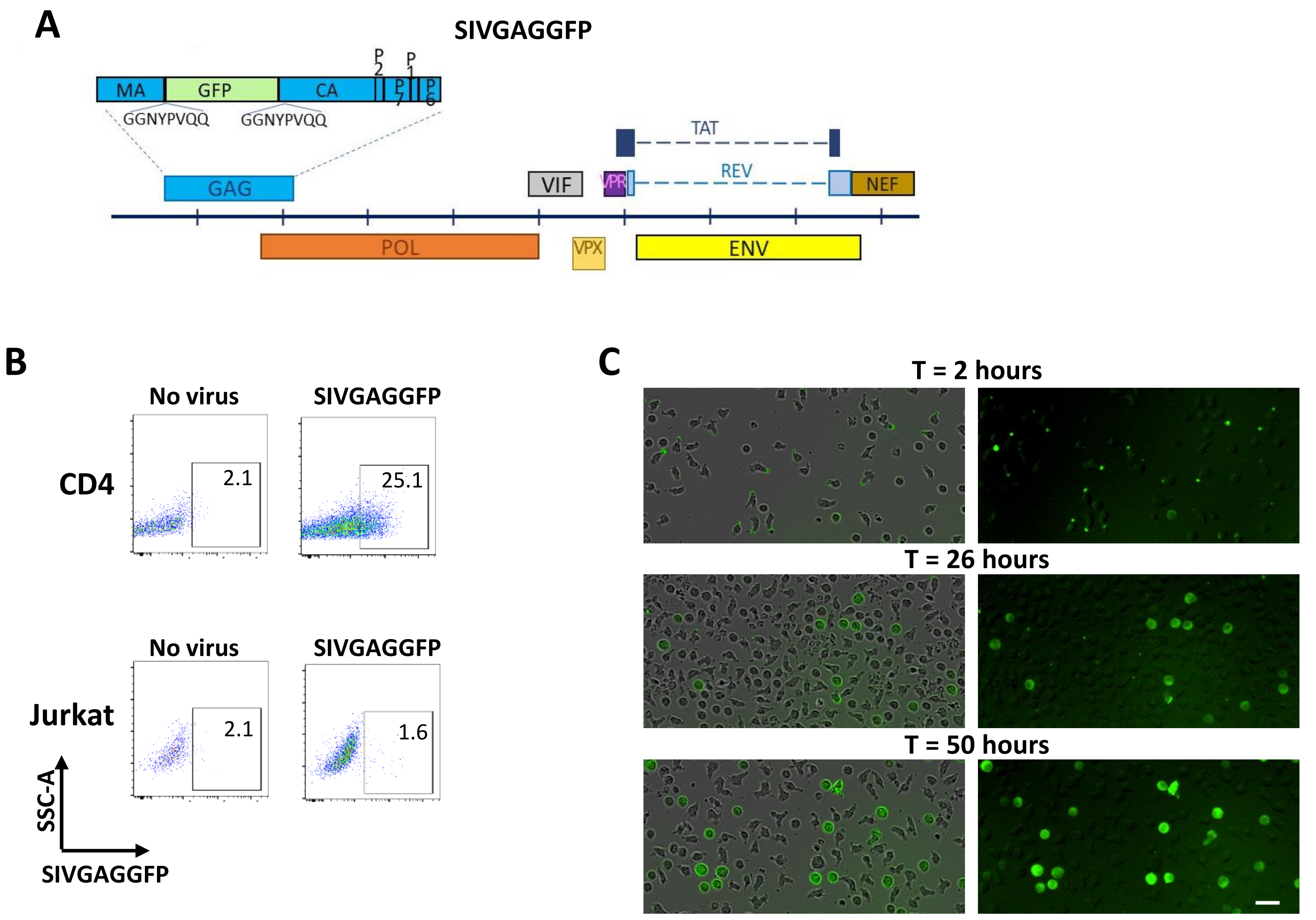


**Supplemental Figure 1. Generation of a fluorescent SIVmac239 virus.** A) The SIVmac239 genome with the GFP coding sequence inserted between the matrix (MA) and capsid (CA) domains of Gag with upstream and downstream GGNYPVQQ protease cleavage sites. B) Flow analysis of SIVGAGGFP binding to RM CD4+ T cells but not to control Jurkat cells. Numbers in plots indicate the percentage of GFP positive cells. C) IncuCyte phase-contrast image superimposed with fluorescent image (left) or fluorescent image (right) of SIVGAGGFP-infected CD4+ T cells after 2, 26 or 50 hours after infection. Localized dots of green fluorescence are observed a few hours after incubation of SIVGAGGFP with CD4+ T cells, while fluorescence was distributed throughout the infected cells 1 or 2 days after infection. Scale bar = 20 µm.
